# Modular Streaming Pipeline of Eye/Head Tracking Data Using Tobii Pro Glasses 3

**DOI:** 10.1101/2022.09.02.506255

**Authors:** Hamed Rahimi Nasrabadi, Jose-Manuel Alonso

## Abstract

Head-mounted tools for eye/head tracking are increasingly used for assessment of visual behavior in navigation, sports, sociology, and neuroeconomics. Here we introduce an open-source python software (TP3Py) for collection and analysis of portable eye/head tracking signals using Tobii Pro Glasses 3. TP3Py’s modular pipeline provides a platform for incorporating user-oriented functionalities and comprehensive data acquisition to accelerate the development in behavioral and tracking research. Tobii Pro Glasses 3 is equipped with embedded cameras viewing the visual scene and the eyes, inertial measurement unit (IMU) sensors, and video-based eye tracker implemented in the accompanying unit. The program establishes a wireless connection to the glasses and, within separate threads, continuously leverages the received data in numerical or string formats accessible for saving, processing, and graphical purposes. Built-in modules for presenting eye, scene, and IMU data to the experimenter have been adapted as well as communicating modules for sending the raw signals to stimulus/task controllers in live fashion. Closed-loop experimental designs are limited due to the 140ms time delay of the system, but this limitation is compensated by the portability of the eye/head tracking. An offline data viewer has been also incorporated to allow more time-consuming computations. Lastly, we demonstrate example recordings involving vestibulo-ocular reflexes, saccadic eye movements, optokinetic responses, or vergence eye movements to highlight the program’s measurement capabilities to address various experimental goals. TP3Py has been tested on Windows with Intel processors, and Ubuntu operating systems with Intel or ARM (Raspberry Pie) architectures.

## Introduction

Throughout the course of evolution, most of visual animals developed eye and head motion to efficiently sample their visual environment. Eye motion frequently involves fast transitions in eye rotation known as saccades, which are common in visual animals with or without high-resolution foveas in their retinas [Kowler, 2011, Land, 1999, Meyer et al., 2020, Wallman and Pettigrew, 1985]. In addition, animals with foveal vision can generate smooth pursuit eye movements when fixating and tracking a moving target. In everyday tasks, humans coordinate their head and eye movements while making voluntary saccadic or pursue eye motions to inspect specific regions of the visual field [Bizzi, 1974, Lanman et al., 1978]. They also generate compensatory eye movements through vestibulo-ocular reflexes (VOR), which are evolutionary well-preserved and neutralize head movements with eye rotations to keep the retinal image stable and eliminate motion blur [Miles and Lisberger, 1981, Raphan and Cohen, 2002]. The optokinetic reflex (OKR) also allows to maintain a stable gaze in a moving environment [Huang and Neuhauss, 2008, Konen et al., 2005].

Head-mounted tracking devices have been used in numerous studies to understand our diverse visual steering strategies within the scopes of sports and navigation [Bahill and LaRitz, 1984, Fegatelli et al., Mann et al., 2007, Shafer et al., 2011], social interactions [Fiedler et al., 2013, Jarodzka et al., 2021, Macdonald and Tatler, 2018, Pfeiffer et al., 2013, Wieser et al., 2009], neuroeconomics [Krucien et al., 2017, Polonio et al., 2015, Reutskaja et al., 2011, Van Der Lans et al., 2008], and cognitive and visual impairments diagnosis [Harezlak and Kasprowski, 2018, Tao et al., 2020, Vidal et al., 2012, Kullmann et al., 2021]. Accommodation, pupil dilation, and convergence further expand the diversity of our visual behavior under different demands of attention, decision making, and depth perception [Fincham and Walton, 1957, Parker, 2007, Preuschoff et al., 2011, Sole Puig et al., 2013, Hoffman et al., 2008, Gibaldi and Banks, 2019].

Within past few decades, software tools for eye tracking advanced substantially with the use of video-based algorithms for gaze estimation [Rufa et al., 2005, Yu and Eizenman, 2004, Cognolato et al., 2018, Lee et al., 2012, Wade and Tatler, 2005, Kim et al., 2004]. This progress occurred in parallel with technological improvements in portable devices and head-mounted displays (HMD), which led to development of more efficient eye and head tracking methods. We chose Tobii Pro Glasses 3 (released in 2020) to perform portable eye tracking, scene capturing, and pupillometry that satisfies our precision criteria for studying naturally-moving visual behavior. Here we introduce TP3Py, an open-source python controller program for live collection and analysis of Tobii Pro Glasses 3 data. Our goal for building the library was to provide a flexible platform for accelerating research in eye/head tracking and behavioral studies. TP3Py allows users to employ custom processing/graphical modules using python programming language. It also allows saving comprehensive behavioral data solely in the host computers including eye movies which are not currently saved within Tobii’s API. The paper is organized into four sections. In section 1, we describe the specifications of Tobii Pro Glasses 3. In section 2, we describe the implementation of TP3Py and built-in modules. In section 3, we explain the calibration procedures. In section 4, we provide a few example recordings and discuss the research directions where the program could have an impact.

## 1 Tobii Pro Glasses 3 Description

Tobii Pro Glasses 3 have similar weight to conventional glasses (77 grams), but are equipped with multiple sensors and cameras to perform eye tracking while freely moving the head (table 1). A HDMI cable transmits the sensor’s output from the glasses to the accompanying recording unit, where video-based eye tracking is performed. The system infers the eye’s direction with 0.6 degrees accuracy by processing the corneal reflection produced by infrared illuminators placed on the glasses (8 per eye). Tobii outputs eye tracking data in a gaze data structure that includes gaze position (gaze2D) in the scene video frame, 3D gaze location for both eyes, and pupil diameters and gaze directions of right or left eyes (table 2). The scene camera’s field of view is 95 and 63 degrees in horizontal and vertical directions. Gaze2D variable outputs (0, 0) or (1, 1) if the subject is looking at the top left corner or bottom right corner of the scene video frame. Tobii is accompanied with rechargeable batteries, each with recording time of 105 minutes.

**Table 1:** Embedded sensors of Tobii Pro Glasses 3

| Sensors | Sampleing frequency | Details | RTP Payload | Encoding format |
| --- | --- | --- | --- | --- |
| Scene Camera | 25Hz | 1920x1080 pixels, YUV420p format | 96 and 97 | H.264 |
| Eye Cameras | 50Hz or 100Hz | Two cameras per eye each<br>256x256 pixels, Single channel | 98 | H.264 |
| Accelerometer | 100Hz | 3 axes | 100 | ASCII |
| Gyroscope | 100Hz | 3 axes | 100 | ASCII |
| Magnetometer | 10Hz | 3 axes | 100 | ASCII |

**Table 2:**
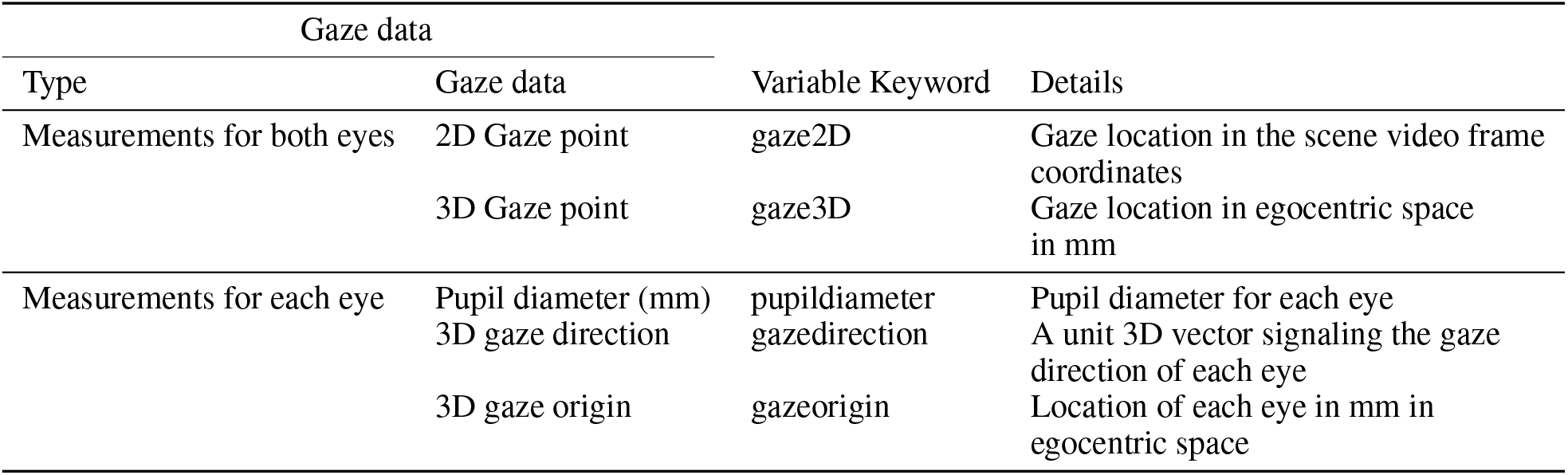
Gaze Data structure

Tobii Pro Glasses 3 transmit multiple digital media packages over wireless via the Real Time Streaming Protocol (RTSP) to the applications. The data packages are specified by the Real-time Transport Protocol (RTP) where distinct data modalities are assigned to a specific payload identifier. The payload identifiers carry information regarding the encoding format, sampling size and sampling frequency of the transmitted data packages. Tobii arranges its data outputs within 6 dynamic payloads from 96 to 101. For instance, scene video frames and scene audio are transmitted over payload 96 and 97. Gaze data structure is transmitted over payload 99 with 100Hz or 50Hz sampling frequency and IMU data is streamed over payload 100. Tobii’s recording unit is also accompanied with a Transistor-Transistor Logic (TTL) circuit which detects a change in the logical value of the TTL input (high to low or low to high) and sends the events over payload 101 that can be used for synchronizing the recording device to stimulus/task controllers.

## 2 TP3Py Program Design

Here we developed an open-source python platform for establishing a flexible communication bridge between Tobii Pro Glasses 3 and client applications. Moreover, the program allows saving comprehensive data including eye videos in the computer host without a need for SD memory cards. We used Gstreamer backend libraries (originally written in C++) to establish the connection and handle the received packages from RTSP server. We also incorporated parallel computing and a Graphical User Interface (GUI) in the TP3Py by using PyQt5 library. TP3Py allows users to develop custom-made processing functions or modules to address a variety of experimental designs and processing objectives.

Figure 1a demonstrates the main interface of the program. Prior to running an experiment, Tobii Glasses should be turned on and connected with WiFi (password TobiiGlasses). The ‘Initials’ and ‘Experiment Name’ text fields correspond to the name/initials of the subject and the conditions of experiment that is being tested. The subject needs to look at the calibration target positioned at 50cm to 100cm distance in front of the eyes after the calibration button is pushed. If calibration is successful, the calibration status becomes green. Calibration can also be performed in the middle of the experiment; however, the data transmission will be temporarily paused. The panel on the right side of the main interface indicates what modules are included in the experiment. Modules access the streaming signals provided by the glasses in live fashion and execute custom processing, graphical, and communicating functions that may be handled within separate threads (fig 1b). For instance, communication modules continuously transfer the acquired signals to other computers where the received information is used for controlling the task conditions. The modules cannot be added at the middle of the experiment. At the end of the experiment, a time period of 5 seconds is needed to make the program ready for the next experiment.

**Figure 1:**
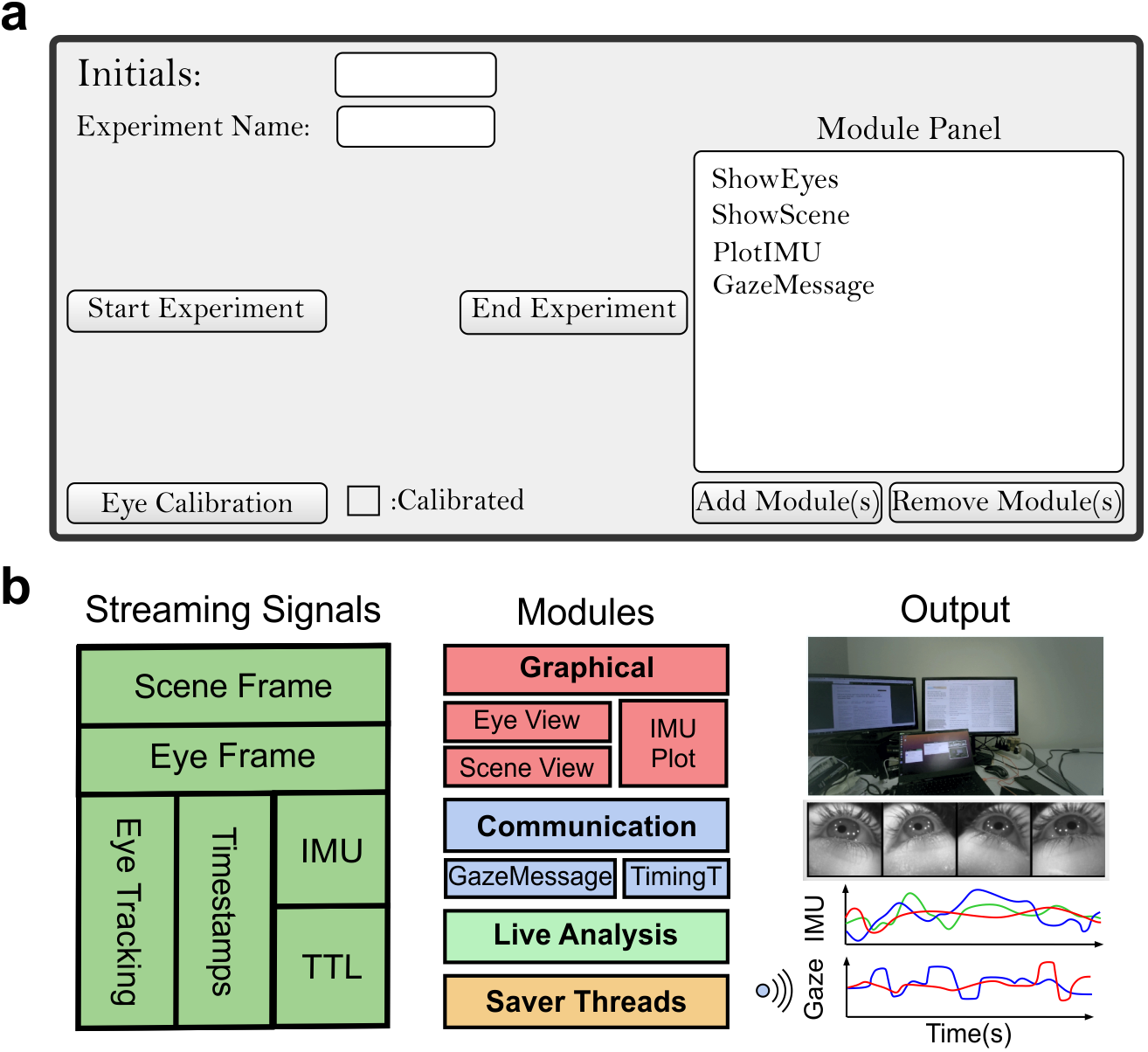
Main Program’s GUI.a. Main graphical interface of the TP3Py including text fields to enter the participant’s initials and experiment name, start/end buttons, eye calibration button and its status (green if calibrated and red if calibration is needed), and module panel for adding and removing modules in the program’s pipeline. **b**. A schematic of TP3Py’s modular pipeline where each module can access distinct data types (i.e., eye frames every 2ms) and execute custom functions for graphical, analysis, and communicative purposes.

### 2.1 Implementing Gstreamer

The main body of Gstreamer contains a pipeline that controls the flow of the application and data pre-processing performed on the received raw packages. We used rtspsrc element at the beginning of the Gstreamer pipeline to connect to the RTSP streaming source. Processing on each RTP packet identified by the payload number are then followed within parallel branches in the pipeline. We used rtph264depay element to extract videos with H.264 format from RTP packets [Wenger et al., 2005] containing the scene and eye video (96 and 98 payloads). Following extraction, we implemented two parallel branches (using tee element) for saving and reading video frames. In the saving passage, the H. 264 formatted videos are directly saved using splitmuxsink element. The splitmuxsink element saves the video in patches of a specified length (default value of a minute) in the .mp4 format. In the reading branch, we employed avdec_h264 element for decoding the video frames and parsed the outcome to the appsink element which makes the received frames accessible to the program in numerical array. Frame’s numerical arrays are then reshaped according to their corresponding image format. For instance, scene images (YUV420p format) are reshaped into a matrix with size of (frame_height*3/2, frame_width). Segmenting the video files in the saving branch makes it easier to load the videos for later usages since the video recorded within the time course of an hour can be immense in size and slow to load. We saved the timestamps of the first video segment frame for time synchronization and aligning the segments. Moreover, all the frame timestamps in the video reading branches are saved.

Appsink element has been also used as an endpoint element of Gstreamer pipeline when processing the rest of the payloads such as IMU data (payload 100), Gaze data (payload 99), or TTL (payload 101) without additional preprocessing. The advantage of using appsink element is that it leverages the raw data in a form of buffers accompanied by the presentation timestamp (PTS) that can be used to synchronize different signals. Video frame buffers are converted to numerical arrays (numpy arrays) while other sources are converted to strings (JSON format) which are later used for saving, processing, and graphical threads. PTS times are generated by a clock that can be set by the Tobii’s internal clock or a local clock of the application host. We will explain the timing characteristics of the system in section 2.5.

### 2.2 Threading, Visualization, and User Interface

We incorporated PyQt5 library for implementing a graphical user interface (GUI) in TP3Py’s pipeline. Not only PyQt5 helped with the front-end visualization but also brought multi-threading functionalities which we used to enhance the performance of the program. Each thread or worker in TP3Py is instantiated from the PyQt5’s QThread class. PyQt5’s most prominent feature is the pyqtSignal and pyqtSlot attributes which allow different threads to communicate. For a connected signal/slot pair, the slot function is called when the signal is emitted, performing a defined function on the received signal. Signals may contain only a digit, number array (numpy ndarray), or characters (string) and they are emitted once a new data acquired or a custom criterion has passed. The Gstreamer thread as an example, continuously uses the pyqtSignals to emit the streamed data to the slots of the other threads that need to carry out saving, processing, or visualizing tasks. Thread classes cannot terminate themselves and they have to send a pyqtSignal to the threads that initialized them for quitting and clearing the memory.

All graphical aspects of the TP3Py are handled with PyQt5 library via the main GUI thread. The main thread manages the user’s input, and it initializes the Gstreamer, saver threads, and other custom-made modules. We utilized PyQt5 widgets for creating text fields, push buttons and other user interfaces in the main window and module interfaces. We further implemented pynput library for inputting keyboard events during the experiment.

### 2.3 Saver Threads

We employed saver threads to continuously save the received data in text files. We connected Gstreamer’s pyqtSignals that contained IMU, Gaze, or timestamps data to the corresponding IMU, Gaze, or time saver-threads. For each experiment all files are saved in a folder named with the experiment’s starting time in the subject name’s directory. All eye video file names are formatted according to ‘eye-ExpName-hour-minute-count’, where ExpName is the input given by the experimenter (figure 1), hour-minute is initial time of the recording, and the count is the number of the video in the segmented sequence (count 0 is the first video). Gazedata, IMU, and video timestamps/TTLs are saved in JSON format in the text file with file names beginning with ‘Gazedata-ExpName-hour-minute.txt’, ‘IMUdata-ExpName-hout-minute.txt’, ‘TSdata-ExpName-hout-minute.txt’. Table 3 presents the variables being saved in each text file.

**Table 3:**
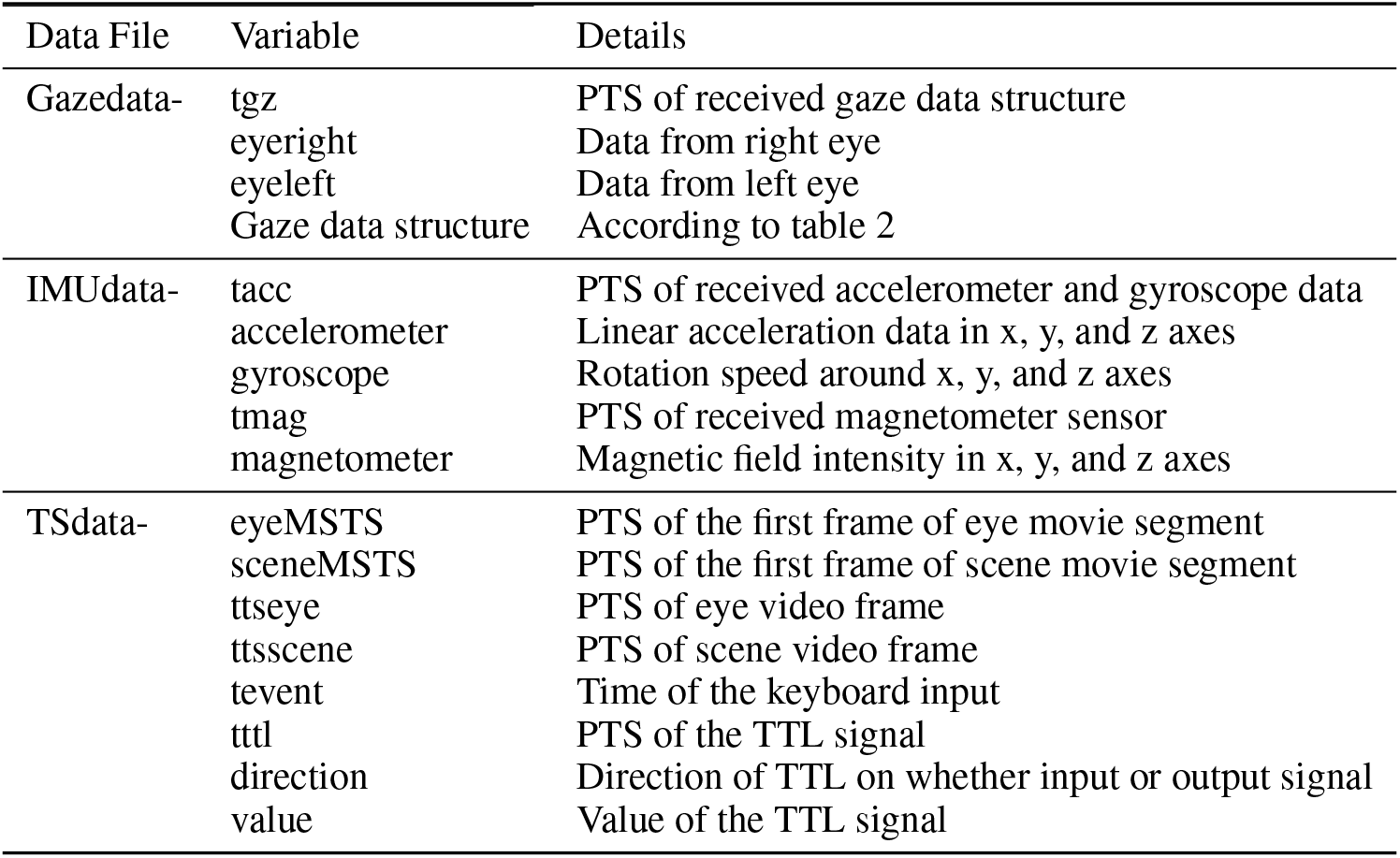
JSON Variables of Gazedata, IMUdata, and TSdata files

### 2.4 Built-in Modules

Custom-made python codes or modules can be incorporated into the TP3Py’s pipeline. The flexibility in accessing Gstreamer signals in addition to the threading capabilities, allow modules to address variety of objectives spanning graphical, analysis, saving, and communicating purposes. Each module is a python class which inherits methods and properties from PyQt5 QObject class and takes the Gstreamer class as an input. Modules can then connect to the Gstreamer signals for accessing the Gstreamer output data such as scene/eye video frames, IMU, and gaze data (fig 1b). Module to module connections can also be established for communicating the processed data if linked properly. Modules are generally organized by a main component and a graphical component. Main components contain the class definition, threading initialization, and callable functions (slots) for processing the received data from other modules/workers. The graphical components characterize the visual interface and layout of the modules. Solely graphical modules generally are not considered as separate threads, since PyQt5 graphics should be rendered in only the main application thread (other graphical libraries such as OpenCV may allow threading without directly using PyQt5’s visualization methods). Nevertheless, Module’s processing loads can be executed within dedicated thread, and this is useful when creating communicating and processing modules.

We incorporated graphical modules for presenting eye videos, scene video and IMU data during the experiment (Figure 1). We converted the color space of the scene (YUV420p) and eye (Gray) movie frames to RGB using opencv library for presenting the video frames into the image canvas of the ShowScene and ShowEyes modules. We also incorporated matlabplotlib library for plotting IMU data in IMUplot module. Furthermore, GazeMessage and TimingTest modules are communicating modules where gaze and TTL data structures are continuously sent to the linked computers over TCP/IP protocol in dedicated threads.

### 2.5 Timing Characteristics

For measuring the timing performance of the program, we employed a closed loop design to measure the time delay between sending and receiving signals. We wrote a Matlab code to send logical (3.3v) triggers using a Data Acquisition Card (NI USB-6001) and 3.5 mm jack adapter connected to Tobii’s recording unit. Upon receiving the triggers, the TP3Py’s TimingTest module sends a character through TCP/IP protocol to the Matlab controller. The controller then finds the time delay between the trigger onset and the time of received character from TP3Py using LAN connection. All the delays in the measurements are averaged across 10 recording sessions each spanning 30 TTL repeats separated within 3 seconds.

The ‘buffer-mode’ variable of the rtspsrc element in the Gstreamer controls the timing conditions. Time reference for data timestamps (PTS) can be synchronized by the sender clock (buffer-mode: 1), internal clock of application host (buffer-mode: 4), or RTP timestamps (buffer-mode: 0). When timestamps are synchronized by the internal application clock, the program is slow (2s). A faster connection is accomplished by synchronizing to the sender clock (0.5 s). However, the timing quality of the data decreases substantially due to interrupted timestamps. The best timing performance is achieved by having the time reference set by RTP timestamps (mean: 140 ms, std: 14ms). RTP timestamps are the timestamps when the data is recorded on the Tobii device, and they differ from the sender clock since Tobii may send data packets at different times irrespective of data acquisition time. Except for the first timestamp, all PTS synchronized to RTP timestamps increase monotonically. Using video modules does not affect the time delay of the program. However, plotting modules such as IMUplot increase the time delay to 180ms. The variations in the timings of the TTL timestamps does not propagate in the relative time differences of TTL’s onsets and offsets (below millisecond accuracy). This suggest that part of the variation in the absolute time measures is inherited in the buffer delay of the DAQ boards connected to the controller computers [Hwang et al., 2019].

All experiments including timing tests are conducted while running TP3Py on laptop (Razor blade, Intel core i7 2.3 GHz, 32 GB RAM) with Ubuntu (20.04) operating system. TP3Py’s open-loop performance has been also tested on windows operating system and portable computing systems with ARM processing architecture (Raspberry Pie 4 model B 8GB RAM). Streaming time delay in Raspberry Pie computing devices are similar to the laptops (150 ms). However, due to the limit in the processing power of the Raspberry, one would need to reduce the scene and eye video frame sizes only in the reading branch of the Gstreamer pipeline without affecting the quality of saved videos.

## 3 Sensor Calibration

Tobii glasses are equipped with inertial and camera sensors that need to be calibrated prior to data analysis. Moreover, the Tobii’s embedded eye tracker requires a single-point calibration procedure for accurate computation of gaze data. Here we provide details on the calibration protocols we performed on the sensors.

### 3.1 InertialSensors

Figure 2a shows the spatial coordinates in which the inertial sensors and eye tracking are based upon. We first calculated the bias and standard deviation of the gyroscope, accelerometer, and magnetometer measurements for each x, y, and z axis while keeping the glasses in a fixed location. The bias and std measurements are later used for estimating head angle using various algorithms. Subtracting average bias from gyroscope data did not result in a Gaussian distribution of the measurements due to the drift present in the gyroscope sensors (figure 2b).

**Figure 2:**
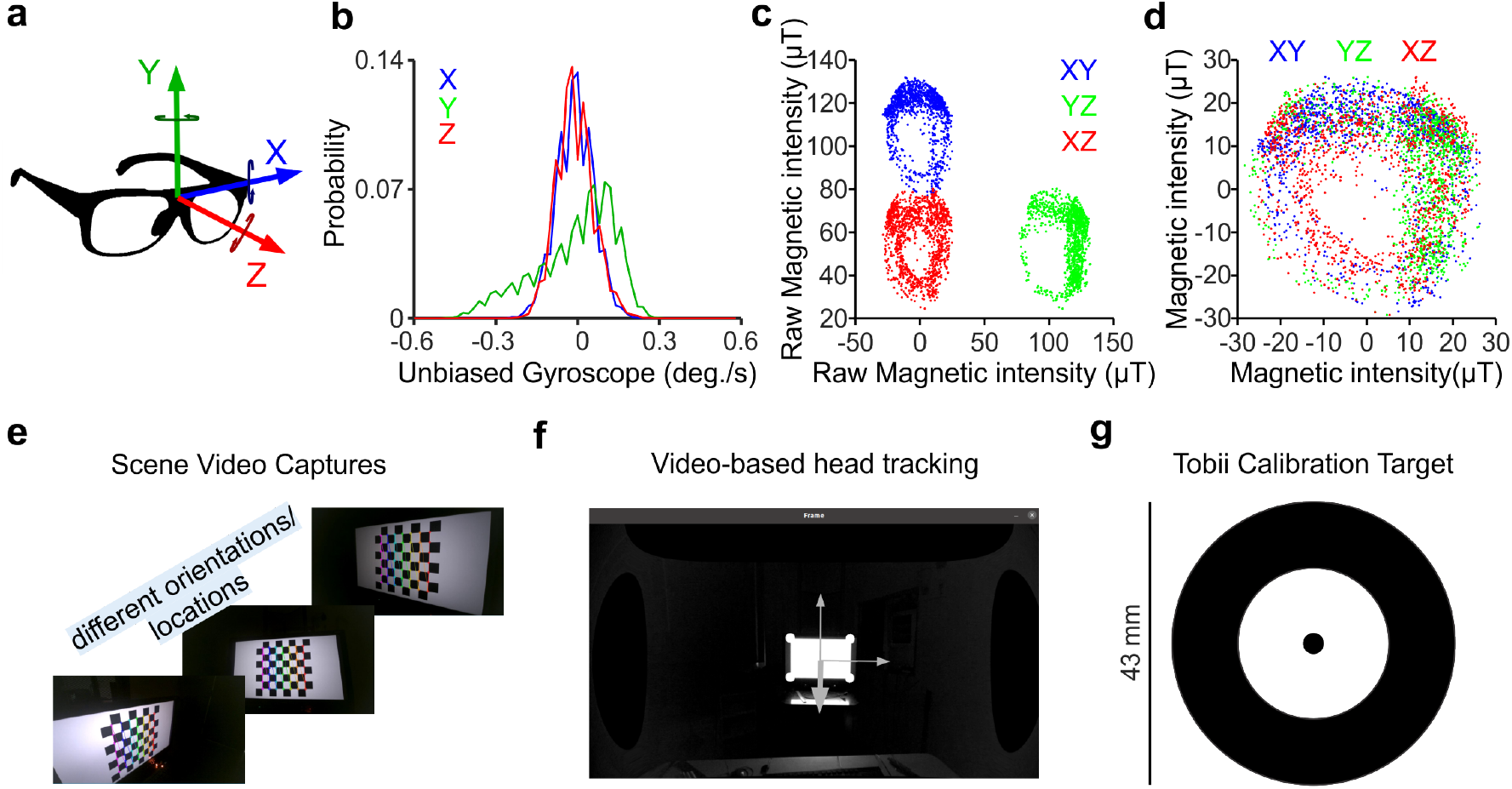
Calibration procedures a. A graph illustrating the spatial coordinates of the glasses. Circular arrows represent gyroscope measurements of rotation speed around the corresponding axes. **b**. Histogram of the unbiased gyroscope measurements along X, Y, and Z axis. **c**. Scatter plot of the raw magnetic field intensity measured along XY, YZ, and XZ axis pairs. **d**. Resulted data scatters after removing hard/soft iron biases from magnetometer sensor. **e**. Schematic of the procedure used for calibrating scene camera by taking snapshots of the checkerboard pattern viewed at different distances and orientations. **f**. Using calibrated scene images to infer the head’s relative position and viewing angle. Solid white circles represent the display corners used for estimating the heading direction. The display coordinates are represented by arrows (the thick arrow represents the Z axis). **g**. Calibration target used in Tobii’s eye tracker. Target background should be white. Inner, middle and outer diameters are 3mm, 23mm, and 43mm correspondingly.

Magnetometers face substantial distortions in their output due to magnetic effects of the surrounding electrical circuit. Permanent magnetic fields in the circuit (hard iron) adds a constant bias to the measurements while other ferromagnetic materials may modulate the magnetic field in particular directions (soft iron). For measuring the hard and soft iron effects on the magnetometer, we slowly rotated the glasses within 8-figure shaped trajectories in various directions. Given the symmetry in the above trajectories, all measurement axis should maintain similar statistics (mean and std). However, due to the hard/soft iron effects, some axes have larger intensity on average than others (figure 2c). In order to counteract these biases, we applied a linear transformation based on the percentile values on each measurement axis to transform the elliptical scatter plots to symmetrical spherical shape centered at zero (figure 2d).

### 3.2 Scene Camera

Spatial relationships between several points in the scene are not preserved in the captured video frames due to the distortions produced by the scene camera optics and sensors. For instance, the length of a bar oriented horizontally or vertically may differ in the captured frames. An intrinsic camera matrix addresses the shift and scale differences in horizontal and vertical directions. Moreover, radial and tangential distortions are produced by the lenses and their displacement relative to the camera sensor. For finding the intrinsic camera matrix and distortion coefficients, one has to calibrate the camera only once.

For camera calibration, we generated 8*8 checkerboard stimulus patterns and captured the scene video of the stimulus from various positions and viewing angles, while slowly moving the camera (figure 2e). Checker’s width and height were set equally to 100 pixels in the display. By knowing the locations of the checker’s intersection points in the display’s coordinates and their corresponding locations in the captured frames, we estimated the calibration coefficients of the scene camera using calibrateCamera function of OpenCV library [Zhang, 2000]. Calibrating the scene camera helps preserve the spatial properties of the scenes which could be used for estimating camera’s location and orientation (figure 2f). In display-based task where the width and height of the monitor is known (the relative locations of the corner points), one could estimate the location and head angle of the observer relative to the display by using the calibrated scene video [Marchand et al., 2016]. Moreover, scene videos provide optic flow measurements that can be also used to infer the head’s direction and its association with the perception [Burlingham and Heeger, 2020].

### 3.3 Eye Tracker Calibration

Tobii’s embedded eye tracker performs a single-target calibration procedure. Upon requesting eye calibration, Tobii detects a unique target (figure 2g) in the scene image and corrects for the eye position biases and drifts. The nominal accuracy of the eye tracking reaches 0.6 degrees. For initializing the calibration, we established a connection to the Tobii API through WebSocket protocol in TP3Py. The calibration target is placed within 50cm to 100cm in front of the observer for 2-4 seconds. The Tobii API then sends a JSON string signaling whether the calibration was successful or not.

## 4 Data Viewers

We developed data viewers in Python and Matlab for viewing and performing post-hoc data analysis. Python data viewer allows previewing the scene and eye videos in a synchronized fashion using the saved timestamps of the scene and eye video frames. We also incorporated image processing functions on the eye images for finding the pupil center and fitting ellipsoids to the pupil shapes that can be utilized for implementing a simple pupil-based eye tracker. Moreover, all data are accompanied by dedicated timestamps which helps to synchronize distinct measurements with high timing accuracy (<1 ms).

### 4.1 Extracting Head/Vergence Angles

We employed sensor fusion algorithms for estimating the head angle using accelerometer, gyroscope, and magnetometer sensors [Guo et al., 2017, Madgwick et al.]. Since the magnetometer sensor is sampled at 10 Hz, we interpolated the measurements to 100Hz to align with inertial sensors. Prior to fusion, all sensors are calibrated using the procedures explained in 3.1. We further measured the vergence angles as follows. Given a unit gaze vector 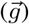 of left or right eye, we calculated the horizontal (*θ*) and vertical (*ϕ*) eye angles using the following equation:

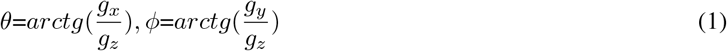

We then calculated the convergence angle of two eyes in horizontal (Θ) and vertical (Φ) planes by subtracting eye angles above.

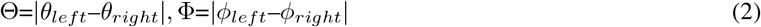

One could also measure the 3D verngence angle (Δ) of the eyes by finding the angle between left and right gaze directions using dot product as below (gaze directions are unit vectors). Locations of the eyes (in the egocentric coordination) are also provided by Tobii which could be used for measuring the viewing distance. However, one could use the z component of the Gaze3D variable which represents the gaze depth in mm.

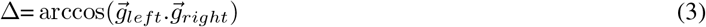

### 4.2 Sample Recordings

We show an unfiltered sample of a recording session in figure 3. In this experiment, the observer fixated on a target placed at 60 cm and rotated his head sequentially in right/left, up/down, and clockwise/counterclockwise directions. Head movements produced counteracting eye movements to minimize the retinal image drift and avoid motion blur. The elicited horizontal VOR eye movements are correlated with the head rotation (r: 0.62, p-value « 0.005) with time delay of 5 ms (using cross-correlation) consistent with other measurements [Ramachandran and Lisberger, 2005]. Furthermore, changes in the head’s pitch angle alters the pupil diameter as well as the horizontal vergence angle signaling an association between VOR and vergence oculomotor afferent signals [Cova and Galiana, 1996].

**Figure 3:**
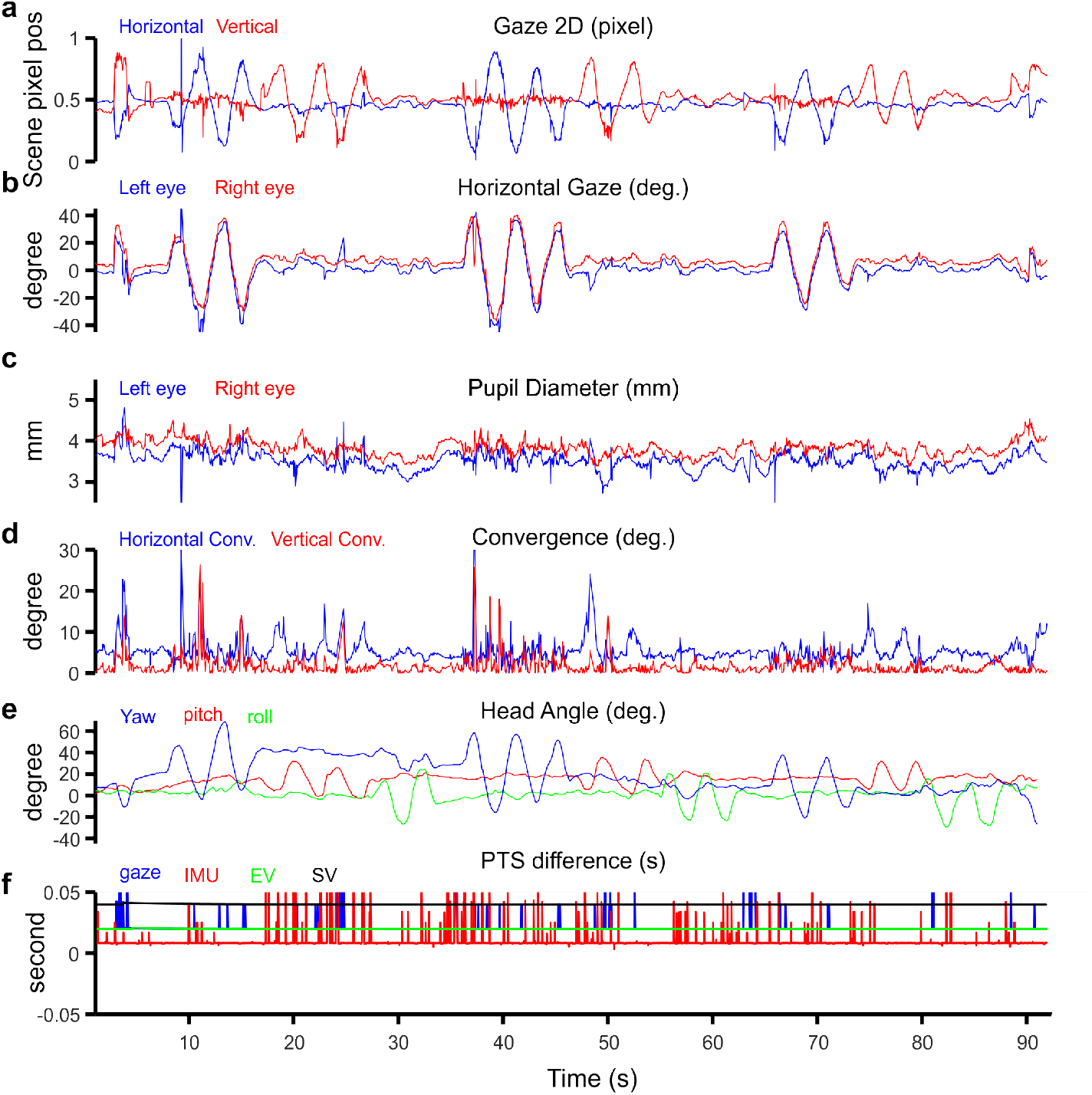
Example VOR recording. a. 2D gaze location of the observer in the scene video plotted across time. **b**. Horizontal gaze angles of left and right eyes over time. **c**. Pupil diameter of left and right eye in mm over time. **d**. Convergence angle of two eyes in horizontal and vertical planes over time. **e**. Head’s pitch, roll, and yaw angles in degrees over time using Fast Kalman algorithm [Guo et al., 2017]. **f**. Presentation Time Stamp differences of distinct streaming media including gaze data (gaze), IMU, scene video (SV), and eye video (EV). Scene and eye videos have the highest timing regularities (0% missed rate) followed by the gaze data with 2% missed rate while IMU data showed the highest timing irregularity (6% missed rate).

We measured timing regularities of the recorded data by accumulating time durations of missed data packages and normalizing by the total experiment time (missed rate). Scene and eye videos showed the highest timing consistency while IMU data showed the lowest. Abrupt head movements and closing the eyes are the main contributors to loosing eye tracking data. It is recommended to adjust the glasses using a proper nose pad for centering the eyes on the eye video frames for better tracking consistency, especially when there are large eye movements.

We further conducted three more recordings, the results of which are presented in the figure 4. In the first recording, the observer had to maintain fixation on a target moving in depth from 10 cm to 100 cm from the observer. The measurements of convergence angle between the two eyes closely follow the convergence eye movement associated with the changes in target location over time. In the next recording (figure 4b), we derive optokinetic responses induced when an observer passively looks at a large (10 deg. size) moving grating stimulus (90 deg. orientation). The pattern of smooth eye movements followed by sudden resetting saccades reproduce the optokinetic nystagmus. In the last recording (figure 4c), we presented two square targets separated by 3 visual degrees horizontally. The subject elicited saccadic eye movements that alternated the gaze location in two targets. Saccadic movements are mediated naturally without imposing any restrictions. Therefore, the results highlights the importance of fast gaze transitions not only when viewing natural scenes but also when assessing simple visual environments [Fabius et al., 2016, Samonds et al., 2018].

**Figure 4:**
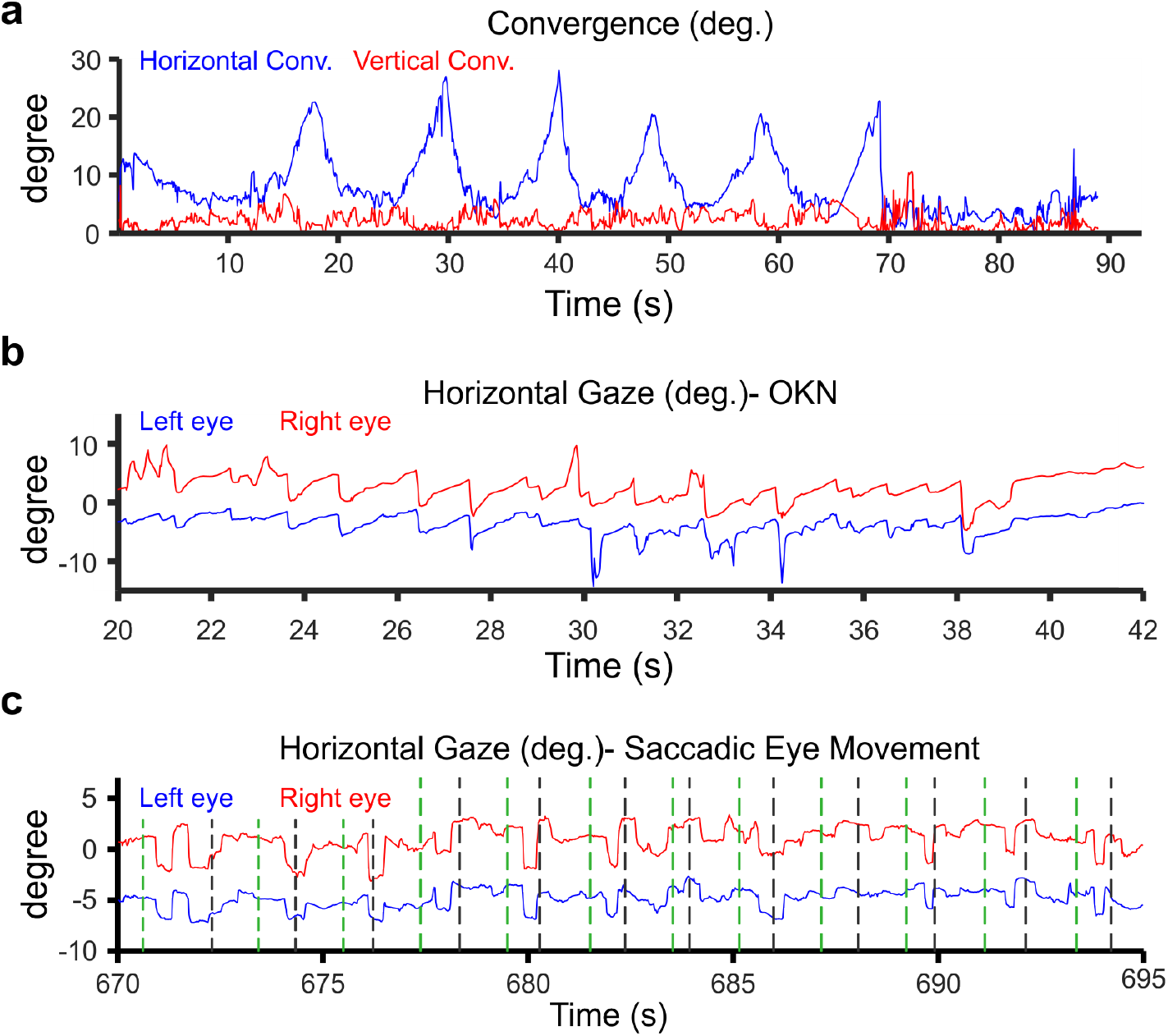
Example recordings of oculomotor vergence, nystagmus, and saccadic motion. a. Horizontal and vertical convergence angle of two eyes over time when an observer fixates on a moving target in depth. **b**. Horizontal gaze angles of left and right eyes over time when observer views a large moving drum stimulus inducing horizontal optokinetic nystagmus. **c**. Horizontal gaze angles of left and right eyes over time when subject is presented with two square targets in the visual field. Green and black vertical dashed lines represent the stimulus onset and offset indicated by switching the TTL’s state. The experiment took 20 minutes to complete, however, time window of 15 seconds has been shown.

## Future Directions and Discussion

We developed a modular controller for acquisition and analysis of multi-modal data including eye/head motion, pupil diameter, convergence, and visual scene content, which helps to provide comprehensive and quantitative measures of visual behavior without imposing restrictions on the head movements. Incorporating TP3Py in open-loop experimental designs provides helpful information on visual behavioral studies. Therefore, the program may help extend the scope of analysis and potential findings when studying psychology, visual perception, and human behavior [Rahimi-Nasrabadi et al., 2021]. The controller’s flexibility and a short time delay (140 ms which is less than a blink duration), allows the program to accommodate various closed-loop experimental designs. TP3Py may be then used for visual rehabilitation and visual training where appropriate feedbacks are required [Trauzettel-Klosinski, 2011, Hernández-Rodríguez et al., 2020] or in studies where visual behavior are mediated in a controlled fashion.

The advancements in portable eye/head tracking devices impact the landscape of freely moving behavioral research. New investigations are needed to fill our current shortfalls on how distinct types of eye/head motions interact in natural setting and what are their consequences for the visual perception [Bouvier et al., 2020, Nau et al., 2018, Shiozaki and Kazama, 2017, Vélez-Fort et al., 2018]. Dynamics of the high-dimensional behavioral motion steered by the natural viewing experiences are more closely addressed when using portable devices [Einhauser et al., 2007, Lappi, 2016, Hayhoe and Ballard, 2005]. As we demonstrated in the example recordings using TP3Py, one can probe the head motion and visual world’s effects on the eye movements, allowing for building more naturalistic image of visual behavior.

TP3Py may also help to accelerate the research in creating novel human-computer interfaces and machine learning tools for extracting relevant information from the visual behavior [Klaib et al., 2021, Kollias et al., 2021, Zemblys et al., 2018, Mildenhall et al., Lee et al., Srivastava et al., 2022]. TP3Py is developed in python programming language which is widely used by the machine learning community and has proved itself efficient for performing artificial intelligence algorithms. Lastly, the software may assist the research in developing video-based eye/head tracking techniques and it may provide benchmarking tools for developing artificial agents that are not only influenced by verbal communications, but also elicit visual and behavioral interactions [Gan et al., Srivastava et al., 2022].

## Acknowledgments

The study was supported by NIH grant EY027361. We thank Vandad Davoodnia for helping to develop the module importation UI.

## Code Availability

The source code of the TP3Py controller and data viewers can be accessed in the link below: Link: https://github.com/SUNYOpt/TP3Py

## Conflict of Interest and Ethics Approval

No potential conflict of interest was reported by the authors. All experiments in human subjects were approved by the institutional review board at the State University of New York College of Optometry and followed the principles outlined in the Declaration of Helsinki. Consent from all participants was obtained prior to the experiments.

## Notes

*Funding* Supported by NIH grant EY027361

https://github.com/SUNYOpt/TP3Py

### Competing Interest Statement

The authors have declared no competing interest.

